# Cutevariant: a GUI-based desktop application to explore genetics variations

**DOI:** 10.1101/2021.02.10.430619

**Authors:** Sacha Schutz, Pierre Marijon, Tristan Montier, Emmanuelle Genin

## Abstract

Cutevariant is a user-friendly GUI based desktop application for genomic research designed to search for variations in DNA samples collected in annotated files and encoded in the Variant Calling Format. The application imports data into a local relational database wherefrom complex filter-queries can be built either from the intuitive GUI or using a Domain Specific Language (DSL). Cutevariant provides more features than any existing applications without compromising on performance. The plugin based architecture provides highly customizable features. Cutevariant is distributed as a multiplatform client-side software under an open source licence and is available at https://github.com/labsquare/Cutevariant. It has been designed from the beginning to be easily adopted by IT-agnostic end-users.

## Introduction

Next-Generation Sequencing (NGS) has opened new opportunities in genomic research such as identification of DNA variations from Genome, Exome or Panel experiments. These data are delivered as files encoded in the standard Variant Calling Format (VCF version 4.0) [1] where the variations are listed together with the genotype information of different samples. Tools such as VEP [2] or SnpSift [3] can be use to add annotations such as genes or functional impact. Biologists can then filter out variants applying customized criteria on these annotations. In medicine, the identification of mutations in rare diseases would be a typical use case. This filtering procedure implements sophisticated software tools that can be easily adopted by end-users who are not necessarily IT-aware.

Several management systems have been developed to ease the usage of the filtering step. GEMINI [4] and VariantTools [5] are command line applications where data from the VCF files are loaded into a relational database managed by SQLite [6]. Filtering can thus be made very efficient using the SQL query syntax. Other tools such as SnpSift [3] or BCFtools [7] apply filters directly while reading the VCF files line by line, thus avoiding the need to create an intermediate data structure. This comes at the cost of poor timing efficiency especially when it is necessary to sort or group variants. While these tools are quite flexible allowing any kind of filtering, the command line interface is not very intuitive, thus reducing the incentive to use it for non IT-specialists.

This called for the development of applications steered by user-friendly Graphical User Interfaces (GUI). Some specializing in diagnostics offer online solutions with a complete set of patient management features but require uploading the VCF files. The most popular of the kind are either private software such as SeqOne [8] and or those distributed under the open source licence such as the recently published VarFish [9]. A major drawbacks of this scheme comes from the transit of a large amount of genetic data through public networks raising on one hand confidentiality and performance issues, and requiring on the other hand a dedicated server which might not be available for every end-users. Moreover, these solutions are tailored for human species data and therefore cannot be adopted for all end-users. GUI Applications that do not require a server and offering an out-of-the-box solution are therefore a preferable solution. The web-based applications VCFMiner [10], BrowseVCF [11] and VCF.Filter [12] implement such a solution. VCFMiner is distributed as a package container running with Docker [13] requiring thus a customized desktop configuration. BrowseVCF provides its own launcher making it quite user friendly but the application is not supported anymore. Both applications import the data from VCF files into an indexed database and provide different GUI forms to create filters. Their main drawback resides in the limited filter settings available through the GUI, complex filters requiring a domain specific language. In addition, web applications offer poor timing performances compared to native desktop applications. Despite the availability of these tools, many biologists still use Microsoft Excel to filter their variants and are facing severe problems [14]. To address the shortcomings of the existing applications, we have developed Cutevariant, a user-friendly and ergonomic desktop application implemented in Python within the Qt5 framework. It takes full advantage of both a GUI and command line user-interface, a Domain Specific Language called VQL allowing the user to build complex filter expressions. It is distributed as a multi-platform client-side software under an open source licence. Thanks to an architecture based on plugins, Cutevariant is fully customizable, allowing to easily extend the application with additional features.

## Materials and methods

### VCF file importation and preprocessing

Cutevariant imports data from VCF files (with or without SnpEff / Vep annotation) into a normalized SQLite database (Figure 1) stored as a *.db file, and optionally with a PED file to describe affected samples and their relationship. Fields from variants and annotations tables are dynamically created according to the content of the VCF file. This importation step proceeds using a VCF parser to produce json-like arrays tailored for populating the SQLite database. It is based on a *strategy design pattern* so that any formats can be supported by subclassing an abstract *Reader* object. The available distribution supports raw VCF files and VCF files annotated with VEP or SnpEff following the ANN specifications [15].

**Fig. 1:**
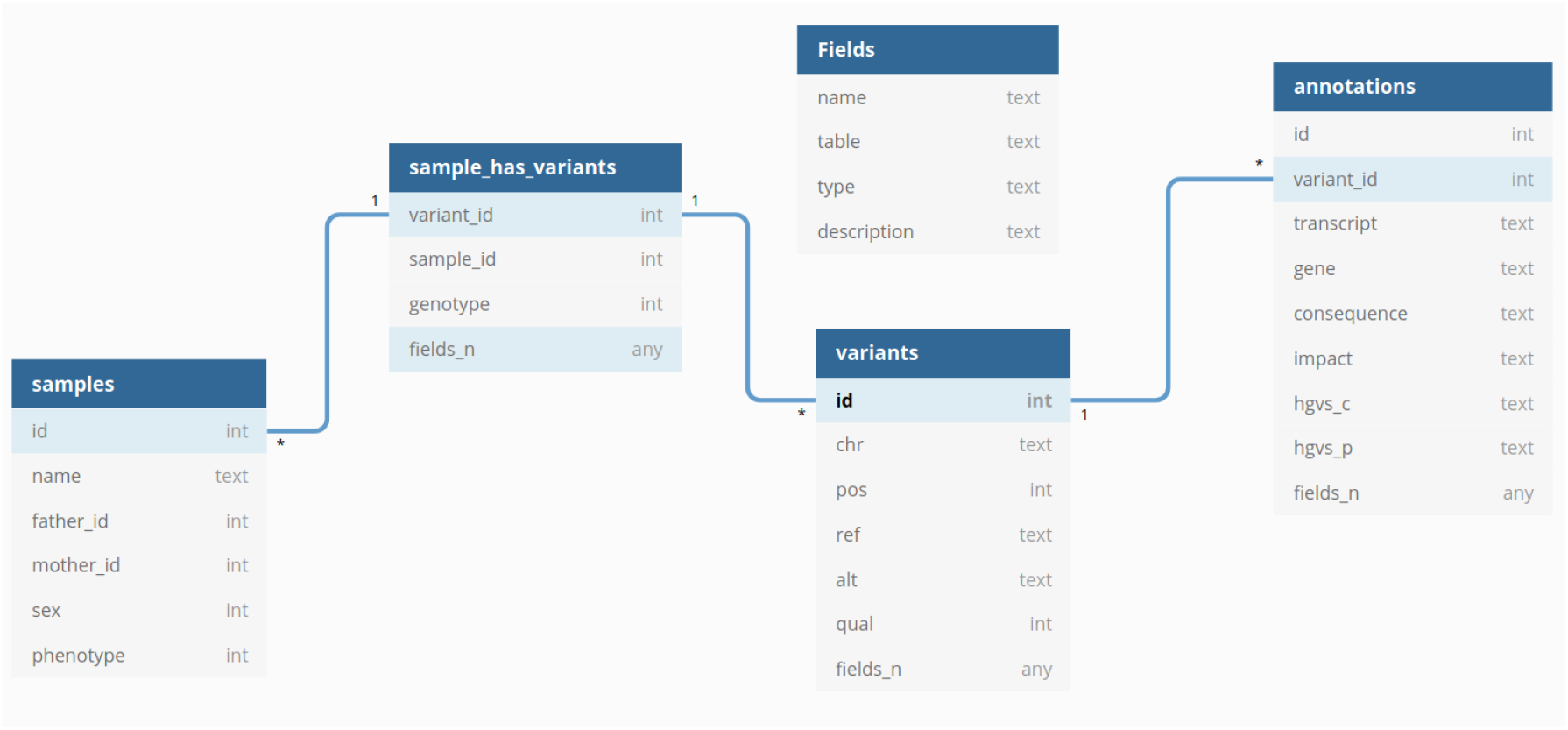
Cutevariant database schema. Only mandatory fields are displayed. *fields_n* are dynamically created during the import step based on the content of the VCF file

Before importation into the database, data are cleaned and normalized following the same procedure as the VT norm [16] application: single lines of multi-allelic variants are split into multiple lines. Computed annotations, not present in the original file, are automatically created. As for example, the *count var* field contains the number of samples that carry the variant. It is thus possible to filter variants present in more than N samples by filtering on this column. This feature is similar to countVar() from the SnpSift [3] filter command.

From the Cutevariant main window, the *new project* button starts a wizard and triggers the importation process. Depending on the size of the input, the importation and indexation process might take some time but this has only minimal impact on the performance since this step is performed only once. Alternatively, VCF files import can be triggered from the command line using the *Cutevariant-cli* button. This feature offers to knowledgeable experts the possibility to integrate the import process at the end of a pipeline.

### User interface layout

The main view (Figure 2) of the Cutevariant GUI displays the list of variants together with their annotations. Several GUI controllers allow the user to update the view and display the list in different formats.

**Fig. 2:**
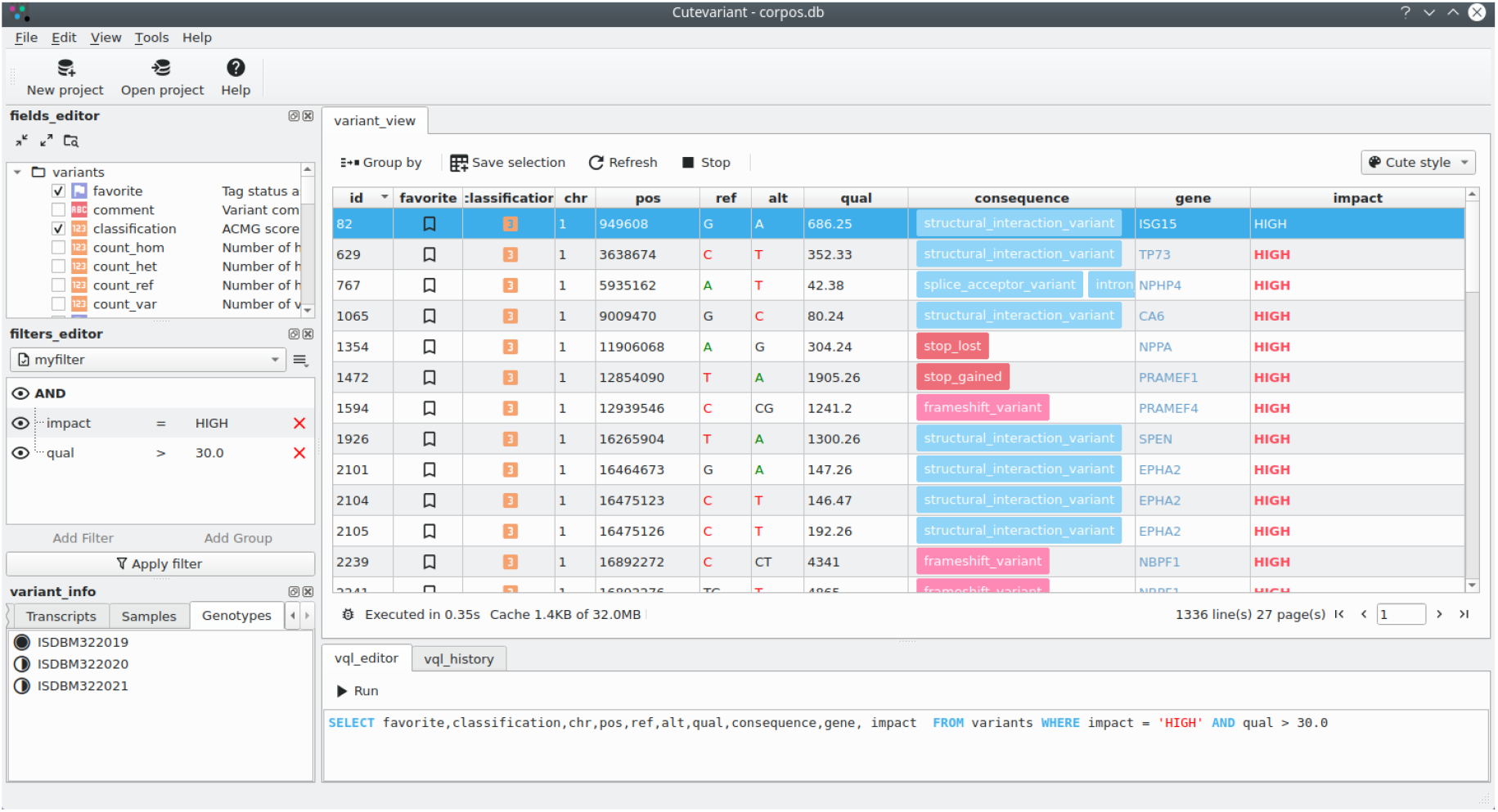
The Cutevariant main view showing the variants list sub-window (middle), different controllers sub-windows but not all are displayed (left) and the VQL editor sub-window (bottom).

- *fields_editor:* to show or hide selected annotations.
- *filter_editor:* to build a nested list of conditional rules with OR/AND binary operators.
- *variant_ info:* to display in an organised way all annotations related to the currently selected variant.
- *source _editor:* to manage different views and perform set operations (union, intersection, difference) and bed file intersections.
- *word _set:* to manage lists of words used to generate simple filters, e.g., filter all variants belonging to a given gene list or a dbSNP list.

Most of these actions end up building a VQL query that can be checked in the VQL-editor sub-window. The variants list can then be updated either with the controllers or by editing the VQL query directly.

### Variant Query Language (VQL)

To facilitate the composition of complex query-filters, the application integrates a Domain Specific Language (DSL) named Variant Query Language (VQL). The syntax of VQL has been designed to look like a subset of the SQL language working on a virtual database schema. It makes use of the Python module textX [17] which provides several tools to define a grammar and create parsers with an Abstract Syntax Tree. VQL queries can be composed in the VQL editor sub-window. However, to avoid forcing users to learn the VQL language, a query can as well be defined from the GUI using the different available controller sub-window listed above. The VQL query is translated through the intermediary of a JSON object into a well formatted SQL query and processed by the SQLite database manager.

As an example, the following VQL query:

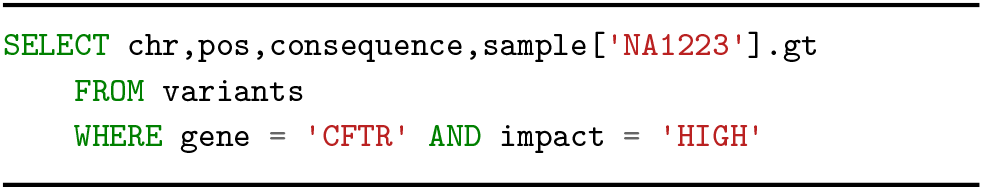

is translated into the following SQL query:

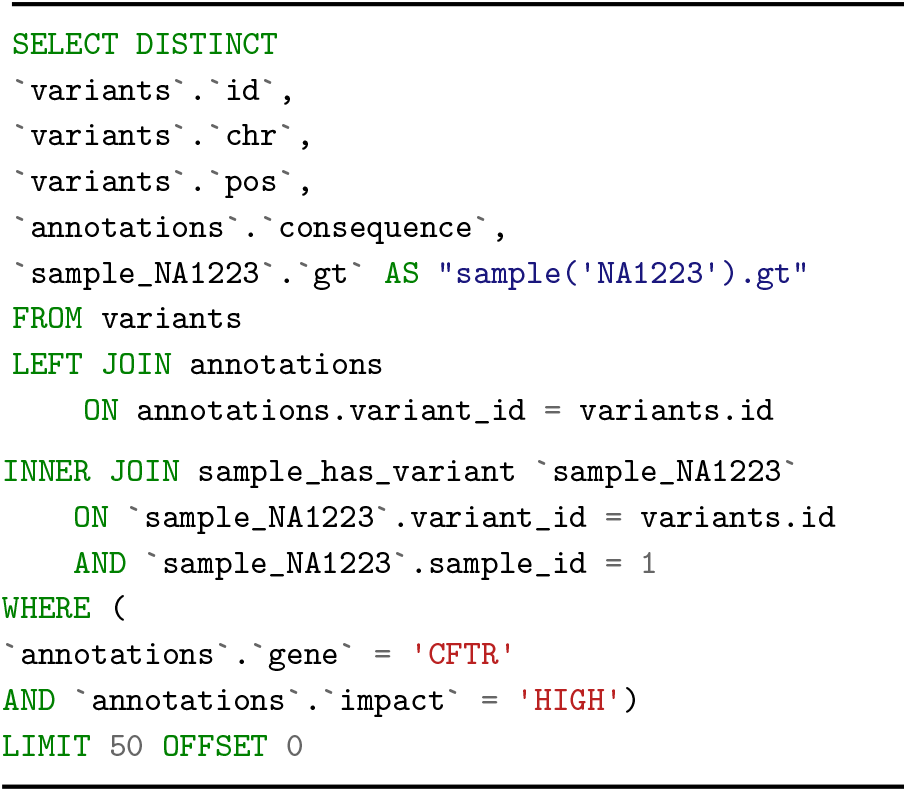

### Filter expressions

Filter expressions are defined from the VQL `WHERE` clause. From the filter editor, it is displayed as a nested set of editable condition rules. Logical (AND/OR) and arithmetic (=, *<, >, ≤, ≥, /*=, IN, NOT IN, IS NULL) operators are supported. Regular expression using the binary ones complement operator (*∼*) and a special *WORDSET* keyword are included as well. This keyword allows the user to test if a fields belongs to a set of words defined a priori. For instance, in VQL, to select all variants from a list of a user-defined genes:

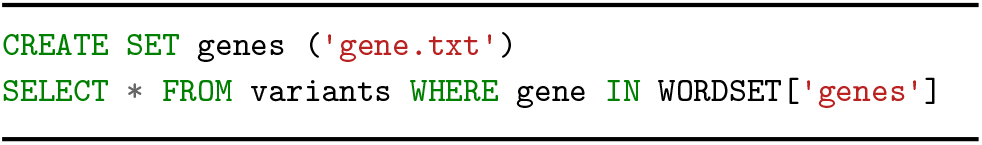

### Group variants

The `GROUP BY` keyword allows the user to split the view in two panels: left the list of groups and right the list of all variants belonging to the selected group. With this feature the exploration is made easier by, for instance, grouping variants by genes helping to detect compound heterozygous.

### Set operation

Just like Variant Tools, Cutevariant supports operations between variant sets. Each query result can be stored in a *view* using the `CREATE` VQL keywords or by clicking the corresponding GUI button. For instance, the following query will create a new view called *new view*.

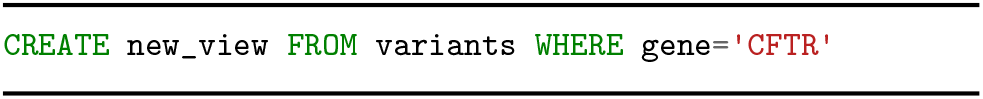

It is then possible to build a query directly from this view. The following query returns the same output as the previous one:

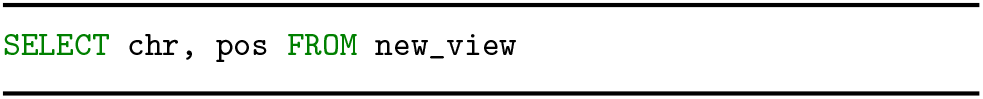

Each view behaves as a set with three operations available (difference, intersection, union) by comparing variants fields on **chr, pos, ref** and **alt**. The following queries show how to create a new view based on different set operation:

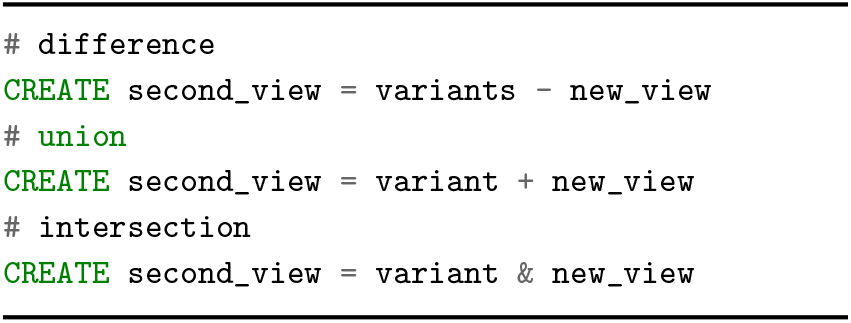

### Plugins architectures

The Cutevariant GUI architecture relies entirely on plugins which source is available in the plugins directory. A plugin consists of a module containing different Python files implementing the creation of a *Plugin class* instance with several overloaded virtual methods. Adding or removing GUI controllers becomes therefore straightforward.

In addition, similarly to excel, cells of the variant view can be formatted conditionally. By subclassing the *Formatter class*, one can change the style of the cell with different colors, text or icons according to the value of the cell. For instance, impact fields with HIGH as value can be displayed with a red background to catch the user’s attention. Currently, Cutevariant supports only one formatters: cuteStyle.

Cutevariant allows the user to build a custom URL from a variant and open it from an external application. This is used for example to open a web link on a dbSNP database or to show BAM alignment from IGV software at the corresponding variant location.

With plugins, experienced users can customize Cutevariant with dedicated features or create new ones and share them with the users community.

### Technical details and continuous integration

Cutevariant is a cross platform application implemented in Python 3.7 using the Qt5 framework for the user interface (PySide2 *≥* 5.11). The VCF parser uses the PyVCF *≥* 0.6.8 library. Syntax and parser of the VQL language rely on the textX *≥* 1.8.0 library. SQLite3 is the database manager interfaced with the Python standard library. The source code and documentation are available on GitHub [18]. Continuous integration are made on GitHub-CI and unit tests are made with the Pytest framework [19]. The application is distributed as windows 32 bits and 64 bits packages. Cutevariant is also available as a Python package from the Python Package Index Pypi [20].

## Results

In Table 1 we list the features available in Cutevariant compared to other applications available on the market.

**Table 1.**
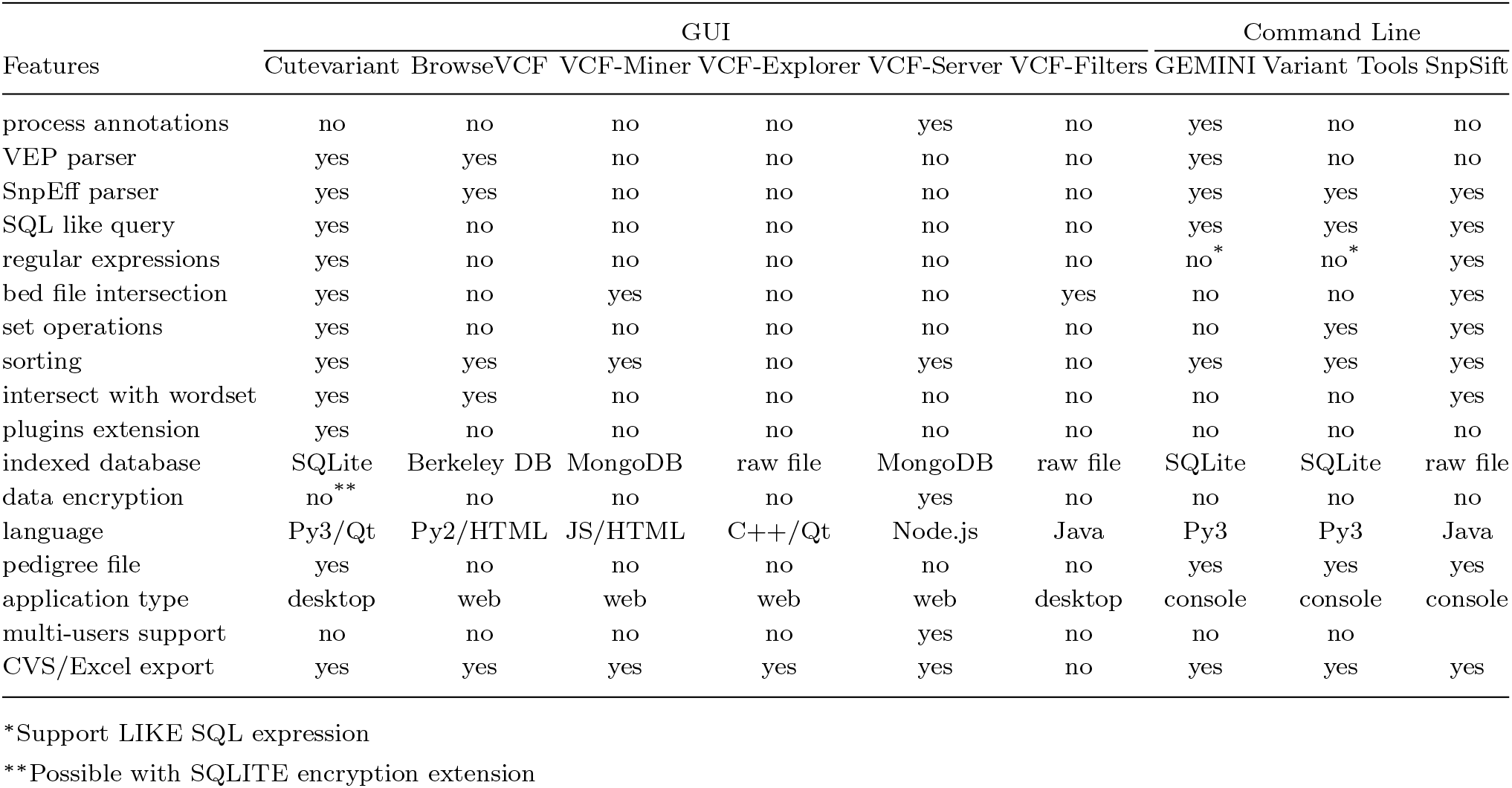
Features available in various applications available on the market.

Cutevariant timing performances for executing importation and query action are reported in Table 2 and compared to the timing performances of VCF-Miner. Other GUI applications could not be tested with our data set for several reasons: parsing error with BrowseVCF, upload size-limit for VCF-Server or lack of indexed database for VCF-Filters and VCF-Explorer. Cutevariant outperforms VCF-Miner except for 1KG.chr22.anno.vcf because of the large number of samples required to compute the joint tables between samples and variants.

**Table 2.** Comparaison of time performance between cutevariant and VCF-miner for importation and query execution. The query used filters variants with *QUAL ≥30 and DEPTH ≥ 30. Executed on Intel(R) Core(TM) i5-3570K CPU @ 3*.*40GHz with 16Gb RAM*

| input file | 1KG.chr22.anno.vcf |  | corpas.quartlet.vcf |  | NA12878.vcf |  |
| --- | --- | --- | --- | --- | --- | --- |
| variant count | 494'328 |  | 300'035 |  | 3'775'119 |  |
| sample count | 1092 |  | 4 |  | 1 |  |
| software | **cutevariant** | **VCF-miner** | **cutevariant** | **VCF-miner** | **cutevariant** | **VCF-miner** |
| importation time | 6600s | 2940s | 78s | 183s | 810s | 2220s |
| query execution time | ≈1s | ≈1s | 0.02s | ≈1s | 0.02s | ≈1s |

### Use case 1: Sars-CoV-2-Analysis

In the context of the Covid-19 pandemia, we have tested Cutevariant to identify mutations along the genome of the Sars-Cov-2 virus. For this, we have downloaded from the ENA database, a dataset (PRJNA673096) with 245 samples stored in a Fastq file produced by the Illumina sequencing plateform using an amplicon librarie. The pipeline is available on github [21].The data originate from the US Delaware Public Health Laboratory. Fastq files have been aligned on the NC045512.2 genome of Sars-CoV-2 with the BWA software [22]. Variants have been called with the FreeBayes application [23] and all 245 samples have been merged into one single VCF file annotated with SnpEff[24]. This file has been imported into

Cutevariant for exploration. We executed a VQL statements (Fig. 4) to extract variants within the gene S and sorted the result by *count var* annotation showing the total number of samples carrying the variant. The sorting process is easily done by clicking on the corresponding header of the view. The mutation p.asp614Gly (highlighted in Fig. 4) is found in 239 samples out of 245. This variant has already been described [25] as a dominant one emerging at the beginning of the pandemia. In the same way, by scrutinizing all the genes, we have identified two others mutation: (ORF1ab)p.Thr265Ile and (ORF3a)p.Gln57His which are exclusive to the North American population [26].

**Fig. 3:**
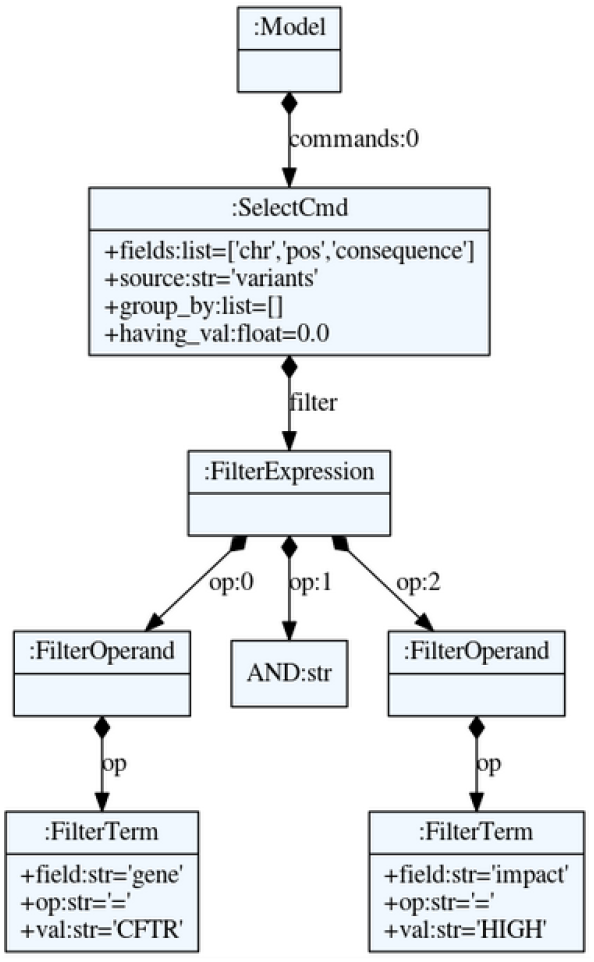
Abstract Syntax Tree (AST) of the VQL query `SELECT` chr,pos,consequence `FROM` variants `WHERE` gene=`‘CFTR’` `AND` impact=‘HIGH’. The AST is parsed into a Python object.

**Fig. 4:**
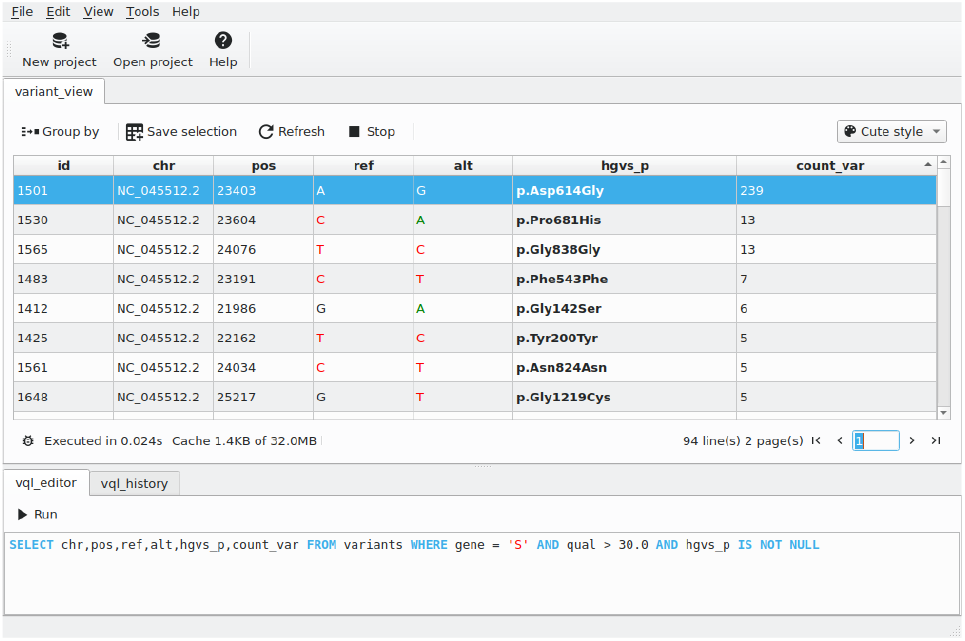
Mutation found in gene S of Sars-Cov-2 by a Cutevariant analysis of 245 samples.

### Use case 2: Cohort analysis

We have repeated with Cutevariant the analysis given as an example by SnpSift [27]. It is a cohort analysis of 17 individuals among which 3 are affected by a nonsense mutation in the CFTR gene (G542*). This analysis cannot be performed with any of the graphics application listed previously (Table 1). After importing the annotated VCF file and the corresponding PED file, the following VQL query was processed by Cutevariant selecting variants with HIGH impact which are homozygous in case samples but are not in control samples. SnpSift uses the following query:

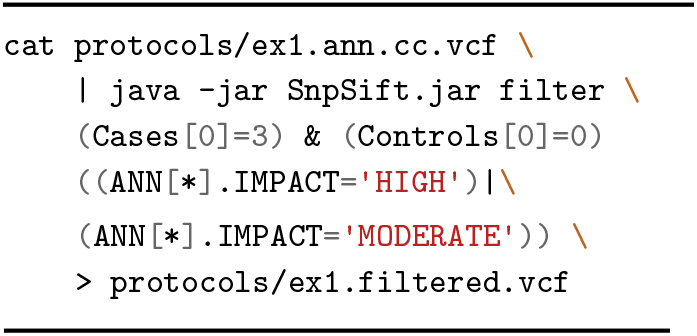

The Cutevariant equivalent VQL query providing the same results reads as:

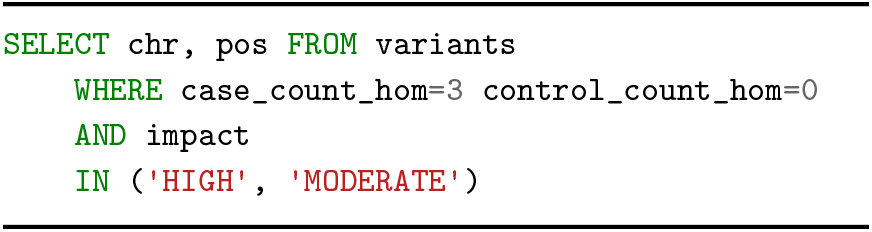

## Discussion

### Performance

Cutevariant is implemented within the open-source Qt for Python [28] that provides a set of Python bindings to build modern user interface. Instead of using native Qt/C++ as coding language, we have opted for Python because it is by far the most frequently used coding language in the bioinformatics community. This choice does not cause any significant performance degradation of the Cutevariant GUI. Execution time for queries performed on a complete genome with many filters can become particularly slow. This long execution time is primarily due to the SQL `COUNT` statement which browses through all the variants to calculate the total number of variants. The table `JOIN` statement is also time consuming. This is the consequence of the choice made for Curevariant, unlike GEMINI, to store samples and a few annotations in separate tables to avoid table denormalization and to minimize disk space occupation. This time penalty has been minimized on one hand by using a memory cache so that identical VQL queries do not need to recalculate the count of variants and, on the other hand, by using asynchronous queries performed in dedicated threads, thus avoiding to freeze the GUI with the progress bar showing the loading status.

### Web app vs Desktop app

Cutevariant is a serverless desktop application and therefore does not provide annotation- or multiuser-features. The annotation step must be carried out upstream at the end of an analysis pipeline by using dedicated tools such as SnpSift or VEP. Multi-users capabilities allow users to share custom annotations and comments. For instance, a user marks a variant as pathogenic and this information is shared among all users. Although this feature is not supported by Cutevariant, it can be delegated to other tools such as MyVariant.info [29]. It provides a database of variants with which Cutevariant can communicate through a REST API. These data can then be used as a source of annotation in the annotation step of the pipeline.

### A general purpose and customizable tool

Cutevariant is a general purpose tool to filter variants and is fully customizable thanks to its plugin-based implementation and thus offers features and modularity that are not available with existing applications. Since Cutevariant is not specific to the analysis of the human genome, it can be use with any VCF file as we demonstrated here with the Sars-Cov-2 example. GUI options dedicated to specific tasks are not hard coded in the application but can easily be added to Cutevariant by creating new plugins. As an example of such added GUI options, the *Trio Analysis* plugin selected from the *Tools* menu users to build from the GUI a VQL filter including transmission mode and the family tree.

## Conclusion

Cutevariant is a new desktop application devoted to explore genetic variations in VCF data provided by next generation sequencing. It is the first GUI software of the kind that integrates both a user friendly graphical user interface and a domain specific language. Starting from a low learning threshold, end-users can easily perform complex filtering to identify variants of interest. Cutevariant is a standalone application that runs on standard desktop computers either under Linux, MacOS or Windows operating systems. The python-based plugins architecture makes the application easily expandable with the addition of new features, thus offering the possibility to involve the biocomputer scientists community at large in new features developments.

## Acknowledgments

We would like to thank Lucas Bourneuf and Pierre Vignet for their contributions.

